# Information-theoretic analysis of a model of CAR-4-1BB-mediated NFκB activation

**DOI:** 10.1101/2023.06.09.544433

**Authors:** Vardges Tserunyan, Stacey Finley

## Abstract

Systems biology utilizes computational approaches to examine an array of biological processes, such as cell signaling, metabolomics and pharmacology. This includes mathematical modeling of CAR T cells, a modality of cancer therapy by which genetically engineered immune cells recognize and combat a cancerous target. While successful against hematologic malignancies, CAR T cells have shown limited success against other cancer types. Thus, more research is needed to understand their mechanisms of action and leverage their full potential. In our work, we set out to apply information theory on a mathematical model of cell signaling of CAR-mediated activation following antigen encounter. First, we estimated channel capacity for CAR-4-1BB-mediated NFκB signal transduction. Next, we evaluated the pathway’s ability to distinguish contrasting “low” and “high” antigen concentration levels, depending on the amount of intrinsic noise. Finally, we assessed the fidelity by which NFκB activation reflects the encountered antigen concentration, depending on the prevalence of antigen-positive targets in tumor population. We found that in most scenarios, fold change in the nuclear concentration of NFκB carries a higher channel capacity for the pathway than NFκB’s absolute response. Additionally, we found that most errors in transducing the antigen signal through the pathway skew towards underestimating the concentration of encountered antigen. Finally, we found that disabling IKKβ deactivation could increase signaling fidelity against targets with antigen-negative cells. Our information-theoretic analysis of signal transduction can provide novel perspectives on biological signaling, as well as enable a more informed path to cell engineering.

## 1. Introduction

Systems biology is an interdisciplinary field that combines experimental and computational approaches to study complex biological systems (Kitano, 2002). It has contributed to the development of mechanistic models to explain biological phenomena and interpret them from novel perspectives. One such perspective has been the application of information theory to biological systems (Uda, 2020). Information theory is the mathematical study of signal transmission and communication (Cover & Thomas, 2012). A central concept of information theory is mutual information. As a measure of dependence between random variables, mutual information carries several advantages. First, unlike correlation, mutual information can be zero if and only if the variables are independent (Shannon, 1948). Second, mutual information makes no assumptions regarding the relationship between two variables. This stands in contrast to various definitions of correlation, which quantify either linear (Pearson’s *r*) or monotonic (Spearman’s *ρ*) relationships and fail to capture more complex patterns (Schober et al., 2018).

One application of mutual information is in signal communication systems, where it is used to analyze relationships between the input signal and the transmitted message. Notably, biological signaling pathways possess properties similar to those of communication systems (Topolewski & Komorowski, 2021). Specifically, a common feature of signaling pathways is to initiate proportionate changes in the concentration of a secondary messenger based on some primary signaling molecule. Viewing extracellular ligands as an input signal and secondary messengers as the pathway’s output, past analyses have treated signal transduction pathways as communication channels and analyzed them from an information-theoretic perspective (Rhee et al., 2012; Waltermann & Klipp, 2011). Such communication channels are commonly described by an inherent property, channel capacity (Uda, 2020). Channel capacity is the highest rate at which information can pass through the channel (Cover & Thomas, 2012). It is achieved by the input distribution that maximizes the mutual information between itself and the corresponding output. Previous work on multiple pathways suggests that this upper limit is typically on the order of 1 bit (Topolewski & Komorowski, 2021), meaning that the pathway’s response can reliably distinguish only 2^1^ = 2 distinct levels of the input signal, i.e., the input’s presence or absence. However, some signaling systems have a better resolving ability. For instance, TRAIL-mediated apoptosis has a population-wide channel capacity of 3.41 bits, meaning that in a given population, TRAIL can specify up to 2^3.41^ ≈ 11 distinct percentages of cells undergoing apoptosis (Suderman et al., 2017). Thus, channel capacity is a useful metric for quantifying the precision by which extracellular ligands can control cellular response via cell signaling.

In recent decades, chimeric antigen receptor (CAR)-engineered cells have emerged as an innovative modality of cancer therapy (Hiltensperger & Krackhardt, 2023). CAR T cells are a type of immune cell genetically engineered to express a receptor comprised of an extracellular recognition domain (specific to the cancer-associated antigen), along with an intracellular signaling domain that actuates cell response after target encounter (Akhoundi et al., 2021). Following infusion into a patient’s bloodstream, CAR T cells can locate and kill cancer cells that express the targeted antigen. Initial designs of the CAR (now termed “first generation”) involved only one signaling domain derived from native T cell receptor and showed little efficacy. However, later work incorporated additional signaling domains originating from immune coreceptors, such as CD28, ICOS and 4-1BB (Srivastava & Riddell, 2015). Termed “second-generation CARs”, these constructs have already shown promising results against certain types of cancer. Excitingly, four of the second-generation CAR T therapies have been approved by the US Food and Drug Administration by 2021 (Sengsayadeth et al., 2021), with the fifth therapy approved in 2022 (Martin et al., 2022). Despite this progress, many limitations remain. For example, existing CAR T therapies are ineffective against solid tumors due to the immunosuppressive tumor microenvironment and low rate of tumor infiltration (Feigl et al., 2023; Guzman et al., 2023). Additionally, CAR T therapy can cause severe side effects, such as cytokine release syndrome and on-target, off-tumor toxicity (Chen et al., 2023). Thus, further research into existing CAR T therapies remains necessary to harness their full potential.

Applying principles of systems biology has yielded success in increasing our quantitative understanding of CAR-based therapies. For instance, one experimentally validated mathematical model has focused on analyzing the pharmacokinetics of CAR T therapy by describing CAR T-mediated cancer cell depletion (Singh et al., 2020). Another study examined signaling processes in CAR T cells by modeling the CAR-CD28 construct’s activation of the ERK pathway (Rohrs et al., 2020). By considering the impact of added CD28 signaling, this study underscored the advantages of second-generation CARs over the first-generation design. The same model was later used to compare the impact of population-wide cell heterogeneity on the activation of CAR T cells (Cess & Finley, 2020; Tserunyan & Finley, 2022b). By applying insights from information theory on CAR-4-1BB constructs, our research group was able to assess the fidelity of CAR-4-1BB-mediated activation of the NFκB pathway by a candidate distribution of the targeted CD19 antigen (Tserunyan & Finley, 2022a). Moreover, it was also possible to evaluate the impact of various perturbations on signaling fidelity. This study produced actionable strategies that can enhance the pathway’s response to encountered targets. However, previous studies focused only on variability caused by the upstream steps in the pathway, without considering variability in the IκBα/NFκB complex the dissociation of which directly leads to nuclear translocation of NFκB (Shih et al., 2011). Furthermore, published works examined a specific example of target antigen distribution without evaluating other biologically-motivated candidate distributions. Hence, further analysis of cell signaling based on insights from information theory can provide a more complete view of CAR-4-1BB-mediated NFκB activation.

In this current work, we set out to develop a deeper information-theoretic perspective of CAR-4-1BB-mediated NFκB activation. By considering cell-to-cell variability in protein concentrations as the source of noise, we have estimated channel capacity of the NFκB pathway in context of the population response. Then, we analyzed the pathway’s diminishing ability to discern contrasting signals with increasing noise levels. Finally, we evaluated the fidelity of NFκB activation when the CAR-4-1BB construct is stimulated with different candidate distributions of targeted antigen. Our work demonstrates that a thorough analysis of signaling systems from an information-theoretic perspective can provide a better understanding of their capabilities. Excitingly, this work, can supply quantitative insights for engineering principles that enable more efficient design in various synthetic biology applications.

## 2. Methods

### 2.1 Network structure

We used our previously published model to analyze CAR-4-1BB-mediated NFκB activation (Fig. 1) (Tserunyan & Finley, 2022a). This deterministic ordinary differential equation-based mechanistic model represents key processes as chemical reactions between various protein species. The model is written in MATLAB and comprises 10 proteins (9 proteins of the pathway and the extracellular antigen), along with 30 kinetic parameters.

**Fig. 1:**
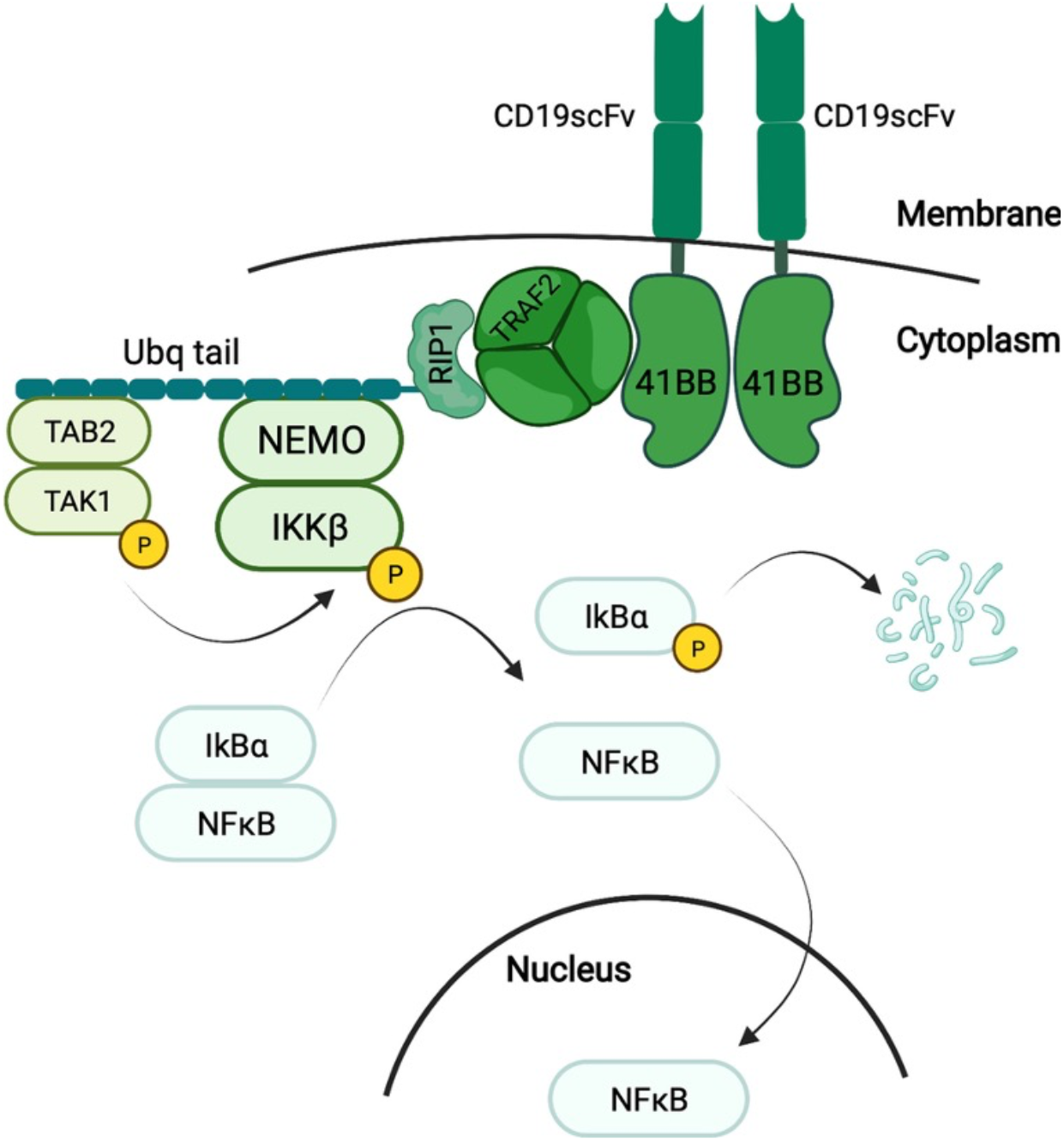
Key steps resulting in CAR-4-1BB-mediated NFκB activation, which manifests in the translocation of NFκB from the cytoplasm to the nucleus (Tserunyan & Finley, 2022a).

Model equations represent the following dynamic process of signal transduction. First, once the extracellular domain of the CAR binds an antigen molecule, the 4-1BB domain of the CAR forms a signalosome, with TRAF2 proteins playing a key role (Zapata et al., 2018). TRAF2 promotes the attachment of K63-ubiquitin chains to RIP1, enabling docking of the TAK and IKK enzymatic complexes (Ea et al., 2006; Kanayama et al., 2004; Lee et al., 2004). After docking, the enzymatic subunit of TAK phosphorylates the β subunit of the IKK complex (Wang et al., 2001). The activation of IKKβ is transitory and typically lasts approximately five minutes due to an auto-dephosphorylating motif near the C-terminus (Delhase et al., 1999). Enzymatically active IKKβ phosphorylates the protein IκBα and initiates its degradation (Zandi et al., 1997). In the resting state, IκBα binds NFκB and sequesters it in the cytoplasm. However, IKKβ-mediated degradation of IκBα releases NFκB from sequestration, allowing it to translocate to the nucleus, where it functions as a transcriptional activator (Ghosh et al., 1998). A negative feedback mechanism ensures that the nuclear concentration of NFκB peaks approximately 30 minutes after activation and then returns to its initial value (Shih et al., 2011). Thus, peak abundance in the nucleus serves as the measure of NFκB activity.

### 2.2 Intrinsic noise of the signaling pathway

We assume that the source of intrinsic noise confounding signal transduction across a population of cells is the variability in concentrations of proteins comprising the pathway. Motivated by the finding that population-wide protein concentrations obey a lognormal distribution (Furusawa et al., 2005), we performed simulations where initial protein concentrations for each CAR cell were independently sampled from a lognormal distribution. We chose the location parameter for the distribution of each protein such that the median concentration across the population would equal the accepted value for that protein. We chose the scale parameter to be the same for all proteins (termed “noise level”), and repeated simulations for multiple candidate noise levels in order to capture general trends in signaling properties. Each simulation regime was performed 2,000 times to ensure a converging sample.

### 2.3 Candidate antigen distributions

We model variability in antigen concentrations encountered by a CAR cell population based on previously published measurements of CD19 antigen (Nerreter et al., 2019). Specifically, fluorescence microscopy showed that the CD19 surface concentration on myeloma cells is in the range of 0.16-5.2 molecules/μm^2^, with antigen-positive cells comprising 10-80% of the myeloma population. Many other cells show CD19 concentration on the order of 0.001 molecules/μm^2^, described as “ultra-low”. Meanwhile, the mean CD19 concentration on leukemia cells from the NALM-6 cell line is approximately 3.4 molecules/μm^2^, with at least 95% of acute lymphoblastic leukemia cells being antigen-positive. Based on this information, we inferred that the surface antigen concentration can be modeled by a bimodal distribution with one component tightly centered around 0.001 molecules/μm^2^ (corresponding to antigen-negative cells) and the other component centered on 3.4 molecules/μm^2^ (corresponding to antigen-positive cells) with a scale parameter of 0.5 to approximate the observed range of measurements. We analyzed four different distributions based on the weight of the positive component: 20%, 50%, 80% and 100% (Fig. S1). Thus, each of these four candidates served as a possible distribution for the input signal (i.e., antigen concentration), and we evaluated the mutual information between a given antigen distribution and the NFκB response it elicited.

### 2.4 Mutual information

Mutual information for continuous variables has been defined via the following equation (Shannon, 1948):

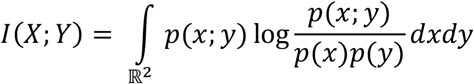

One approach for computing the mutual information of two continuous variables based on a statistical sample is to discretize them (Ross, 2014). However, this can cause large discrepancies in the calculated number based on the choice of discretization scheme. As an alternative, we have used the Kraskov algorithm to compute mutual information, which avoids explicit discretization (Kraskov et al., 2004). Instead, it relies on a *K*-nearest neighbors procedure and tends to deliver more consistent results.

### 2.5 Channel capacity

The long-standing Blahut-Arimoto algorithm and its variations have been used for computing channel capacity for discrete variables (Cover & Thomas, 2012). Unfortunately, an exact algorithm for transmission systems where both the input (e.g., encountered antigen concentration) and the output (e.g., nuclear concentration of NFκB) are continuous, has not been published to our knowledge. While efforts have been made to overcome this limitation, those attempts incorporated additional assumptions regarding the structure of probability distributions and the relationship between input and output, and the calculated channel capacity can fluctuate if those assumptions are not satisfied (Jetka et al., 2019).

To approach this problem, we have devised an iterative heuristic procedure to estimate the channel capacity for CAR-4-1BB-induced NFκB activation. To start, we recalled the equivalence, which expresses mutual information as the difference between entropies of a variable’s prior and posterior distributions (conditioned on the second variable) (Cover & Thomas, 2012; Rhee et al., 2012):

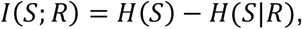

where *I(S; R)* is the mutual information between random variables *S* and *R* (denoting the input signal and output response, respectively), *H(S)* is the entropy of *S*’s prior distribution and *H(S*|*R)* is the conditional entropy of *S*’s distribution provided *R*, with all three quantities lower bounded by zero (Uda, 2020). Since channel capacity is the maximum mutual information that can pass through the channel, we looked for an input signal distribution that has a large prior entropy, but a near-zero conditional entropy.

Next, we recalled that channel capacity is interpreted as the logarithm of the largest number of distinct input levels resolvable by the channel (Topolewski & Komorowski, 2021). For example, if channel capacity is *n* bits, then the system can resolve up to 2^*n*^ distinct input concentrations. It follows that this capacity could be achieved by an antigen distribution if it meets two conditions. First, in the antigen’s prior distribution, these *n* distinct levels should be equally probable (to maximize *H(S)*). Second, by knowing the secondary messenger’s concentration *R*, we should be able to infer with negligible error which of the *n* concentration levels was used for antigen (to minimize *H(S*|*R)*). In the context of NFκB signaling, which shows a sigmoidal dose response, we can choose specific antigen levels that would be as easy to resolve as possible. For example, if we are trying to distinguish between two antigen concentrations, then the most resolvable pair would be the near-absent and saturating values, since activation caused by near-absent antigen would be indistinguishable from basal levels, while activation caused by the saturating concentration would be higher than for any intermediate value. Similarly, the set of three antigen concentrations where the respective outputs are easiest to distinguish would be those, which cause 0%, 50% and 100% activation in the system, since the response distribution corresponding to the added “middle” value would be equidistant from the other two and show least overlap with them (assuming response distributions are symmetrical). To generalize, if we have *n* distinct antigen concentrations, they would be most resolvable if the activation levels they elicit are evenly spaced 100%/(*n*-1) apart. Such concentration values can be found with the dose response curve.

Given this assessment, we proceeded as follows. First, we assumed that the pathway encodes only two distinct antigen concentrations, “low” at 0.7 molecules/μm^2^ and “high” at 70 molecules/μm^2^. This was motivated by the fact that at 0.7 molecules/μm^2^, the pathway shows virtually no activation, and nuclear NFκB remains at the basal level (Tserunyan & Finley, 2022a). Meanwhile, at 70 molecules/μm^2^, the pathway is fully saturated, and further increases in the antigen level show no appreciable increase in the concentration of nuclear NFκB. We obtained the response distribution for this setup and computed the mutual information between concentrations of antigen and nuclear NFκB. If this value was close to the entropy of the input distribution (implying *H(S*|*R)* = 0, i.e., absolute certainty of antigen concentration provided the NFκB response), we concluded that the pathway could resolve additional input levels. Hence, we increased the number of distinct antigen concentrations by one and continued this process until the mutual information between antigen concentration and NFκB response stops increasing. Following this procedure, we took the maximum mutual information observed for each noise level as corresponding channel capacity (Fig. S2). When disabling IKKβ deactivation, the dose response curve shifts, necessitating a different choice of candidate antigen concentrations, but the procedure is otherwise the same (Tserunyan & Finley, 2022a). In this case, we used 0.1 molecules/μm^2^ and 20 molecules/μm^2^ as the starting “low” and “high” antigen concentrations.

The fidelity of NFκB activation can be confounded by two sources that diminish the mutual information between antigen concentration and nuclear NFκB. One is the stochasticity in the cellular abundance of NFκB and IκBα, the dissociation of which directly leads to NFκB’s nuclear translocation. Another source is the variability in upstream proteins that lead to IκBα’s phosphorylation and degradation. To discern how each of these sources affects channel capacity of the NFκB pathway, we performed Monte Carlo simulations for two scenarios: first, where upstream proteins are variable, but the concentrations of NFκB and IκBα are fixed to the same value for each simulation; and second, where NFκB and IκBα are variable along with all the other proteins. For the rest of the simulations, we focused solely on the more biologically realistic scenario with all proteins being variable (including NFκB and IκBα). Finally, since past experimental research has shown that the fold increase in NFκB nuclear concentration at its peak can be more informative than the absolute value (R. E. C. Lee et al., 2014), we repeated our calculations of channel capacity by taking fold increase in nuclear NFκB following stimulation, relative to baseline, as the output variable.

### 2.6 Assessing error rates

To demonstrate that decreasing channel capacity hinders the pathway’s ability to discern antigen concentrations, we examined how its response can distinguish between “low” (0.7 molecules/μm^2^) and “high” (70 molecules/μm^2^) levels of antigen (Fig. S3). First, we obtained distributions of peak nuclear concentration of NFκB for each of those antigen levels. Then, we devised a backwards procedure by noting whether the likelihood of each obtained absolute response value was higher under the assumption of coming from the “low”- or “high”-stimulated response distribution. When a specific value of NFκB concentration came from simulations with a “high” antigen concentration but its likelihood is greater under stimulation with “low” antigen concentration, we considered this an instance of a “false negative”. We defined the converse case as “false positive”. We represented these data as percent of false positive and false negative estimates for each noise level. Finally, we repeated this procedure by using fold change in nuclear NFκB as response variable.

Given the difference between dose response curves for the unperturbed system and with disabled deactivation of IKKβ, we performed the analysis for the second case by accepting 0.1 molecules/μm^2^ and 20 molecules/μm^2^ as the “low” and “high” concentrations.

## 3. Results

### 3.1 Channel Capacities

First, we computed channel capacity implied by our model for CAR-4-1BB-mediated NFκB signaling. We evaluated channel capacity for two simulation conditions: (1) varying concentrations only for proteins upstream of the NFκB/IκBα complex (IκBα and NFκB concentrations are fixed); and (2) varying concentrations for all proteins, including IκBα and NFκB. For each condition, we focused on two response measures that characterize the amount of information carried about antigen concentration by NFκB: peak value of NFκB concentration in the nucleus following stimulation, and the peak fold change in nuclear NFκB relative to baseline.

We observed that with all other factors fixed, channel capacity decreases with increasing intrinsic noise levels (Fig. 2a). However, results from various simulation conditions differed in the extent of this decrease. When the concentrations of NFκB and IκBα were fixed across the population, channel capacity of the signaling system was the same whether we considered the absolute response of nuclear NFκB or the fold change (Fig. 2a, blue and yellow). This is anticipated since a fixed pre-activation concentration of NFκB and IκBα means that fold change is identical to the absolute concentration up to a constant multiplier. However, with variable concentrations of NFκB and IκBα, channel capacity entailed by the absolute response decreased by approximately 1 bit for all tested noise levels (Fig. 2a, green). This decrease in channel capacity corresponds to a 50% reduction in the number of distinct antigen concentrations the pathway can resolve due to variability in NFκB and IκBα levels alone.

**Fig. 2:**
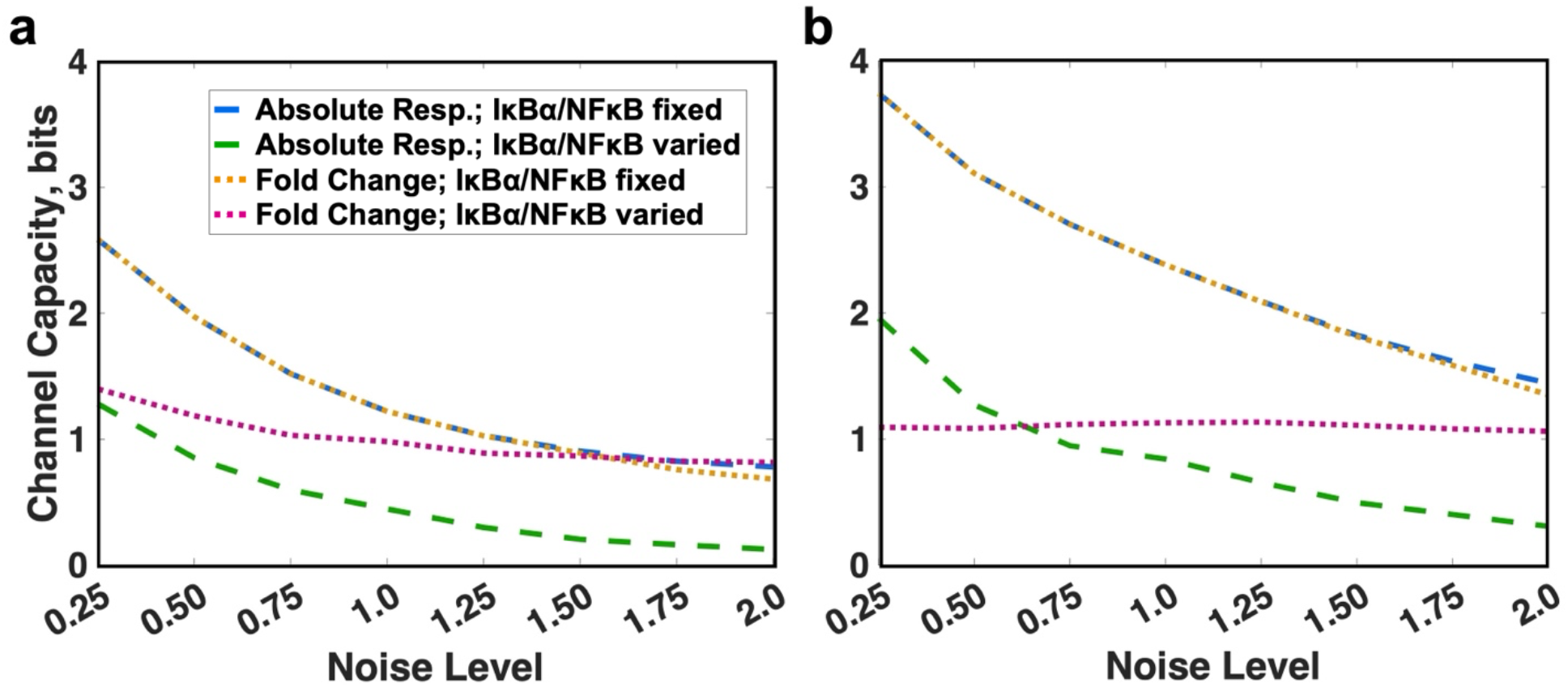
Channel capacity of CAR-4-1-BB-mediated NFκB activation calculated for different simulation conditions either (a) for the default parameters or (b) disabled IKKβ deactivation. In each case, channel capacity was computed for the nuclear concentration of NFκB in a population of cells with fixed (blue) or variable (green) pools of IκBα and NFkB, in addition to the channel capacity entailed by the fold change with fixed (yellow) or variable (magenta) pools of IκBα and NFkB.

When the abundances of NFκB and IκBα vary, channel capacity entailed by fold change of nuclear NFκB proved to be comparatively consistent across noise levels (in the range of 0.82-1.4 bits) (Fig. 2a, magenta) and higher than channel capacity of the absolute response (Fig. 2a, green). This difference grew from 0.12 bits for noise level 0.25 to 0.70 bits for noise level 2.0. This suggests that when overall noise level was relatively low, most variability in activation levels was attributable to stochasticity in upstream signaling species, so that measuring fold change did not result in a significant information gain. In contrast, at higher noise levels, response variability can be mostly attributed to the stochasticity in cellular levels of NFκB since the fold change of nuclear NFκB showed significant information gain over the absolute response.

In our previous work, we found that disabling the deactivation mechanism of IKKβ could increase the mutual information between the distribution of antigen concentration and NFκB activation (Tserunyan & Finley, 2022a). Now, we aim to verify whether this observation is limited to a specific input distribution or reflects a change in transmission abilities of signaling system. Thus, we carried out similar calculations of channel capacity with disabled deactivation of IKKβ (Fig. 2b). With fixed IκBα and NFκB concentrations, we observed a consistent increase in channel capacity by approximately 1 bit in the perturbed model for all noise levels. This was true for both absolute response and fold change (yellow and blue in Fig. 2b compared to Fig. 2a).

When IκBα and NFκB vary, disabling IKKβ deactivation resulted in an increase in channel capacity of the absolute response by up to 0.7 bits (green in Fig. 2b compared to Fig. 2a). Interestingly, channel capacity of the fold change is predicted to be almost independent of the intrinsic noise parameter and confined to 1.06-1.13 bits (Fig. 2b, magenta). Notably, for low noise levels, fold change had a smaller channel capacity than absolute response. However, this trend did not hold at noise levels greater than 0.5, when the capacity of the absolute response decreased with added noise, while the capacity of fold change stayed the same.

In summary, we found that when IκBα and NFκB concentrations were fixed, the NFκB pathway’s capacity to transmit information about the input signal was independent of the measure of signal transduction (absolute response vs. fold change of NFκB). This remained true when IKKβ deactivation was disabled. If IκBα and NFκB concentrations were varied, the unperturbed pathway had a higher capacity to transmit information via the fold change of NFκB. If IKKβ deactivation was disabled, the pathway could better relay information via the fold change of NFκB only at higher noise levels.

### 3.2 Signal discernibility

To further understand how intrinsic noise affects signal transmission for CAR-4-1BB-mediated NFκB activation, we decided to stimulate the pathway *in silico* with two contrasting antigen levels while varying all protein concentrations. Then, we evaluated how effectively NFκB activation can distinguish those signals. We did this by observing whether a given value of NFκB response obtained from “high” stimulation is more characteristic of the “low”-stimulated regime (“false negative”) and vice versa (“false positive”). Following this approach, we saw that for all noise levels, the false positive rate remains near 0% for both the absolute response and the fold change (Fig. 3a, blue and yellow). In contrast, the false negative rate rose to 80% for absolute response depending on the level of intrinsic noise (Fig. 3a, teal and purple). Notably, the false negative rate was 20-25% lower if the fold change of NFκB is considered instead. This means that the cells are predicted to transmit information more accurately via the fold change of NFκB when contrasting stimuli are provided.

**Fig. 3:**
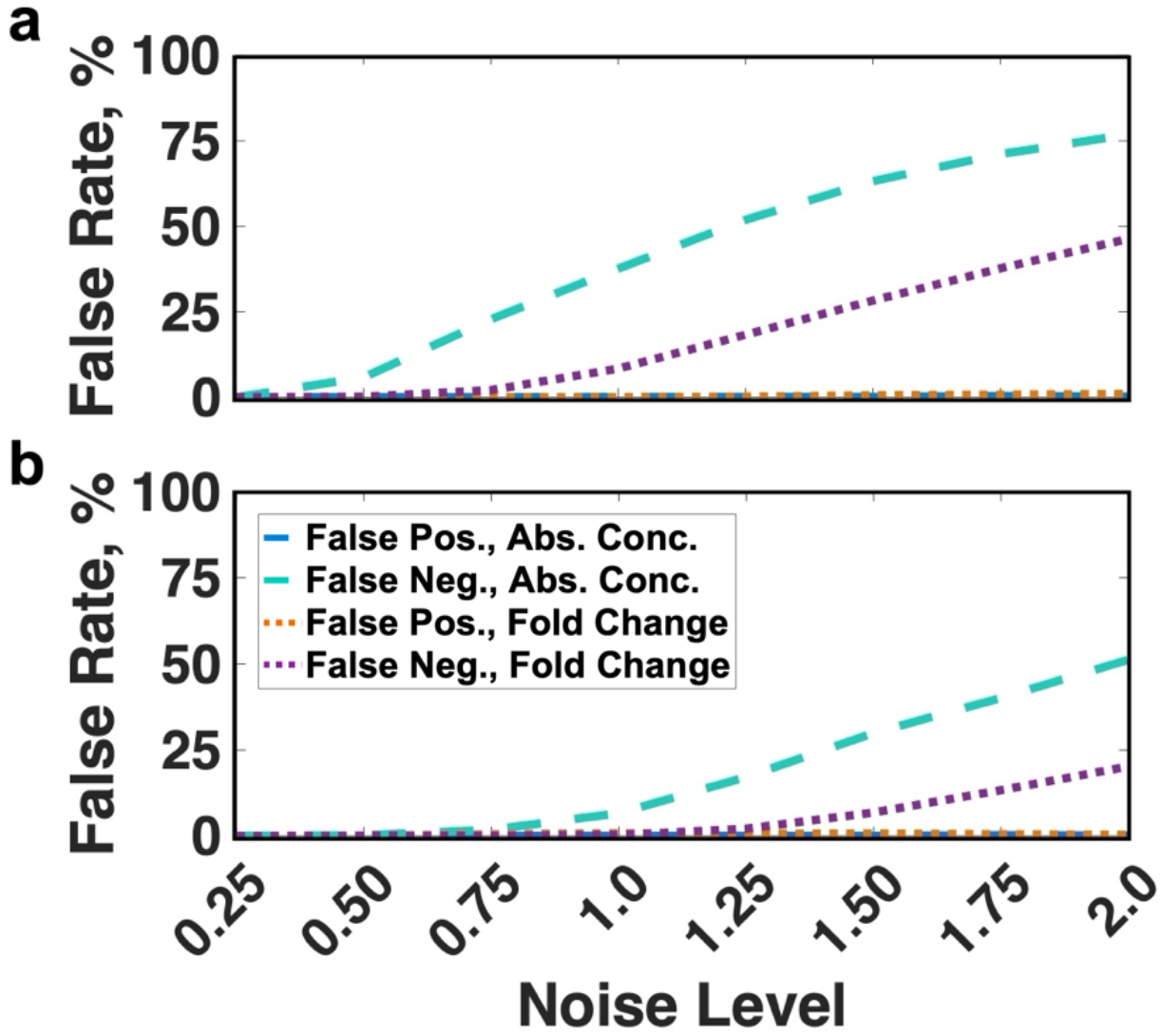
Ability of correctly distinguishing contrasting antigen concentrations based on NFκB activation either (a) for the unperturbed model or (b) disabled IKKβ deactivation. Error rates were computed in a population of cells based on absolute response (blue, false positives; teal, false negatives) or fold change (orange, false positives; purple, false negatives).

We performed the same analysis of the error rates while disabling IKKβ deactivation. Here, we observed that while the false positive rate remains at zero, the false negative rate is up to 35% lower compared to the unperturbed system (Fig. 3b). Notably, as with the unperturbed model, cells were better able to transmit information about the input signal via the fold change in NFκB than the absolute response (Fig. 3b, purple is lower than teal).

These findings demonstrate an interesting agreement with our estimate of the channel capacity. Errors in transmitting two contrasting inputs would imply a channel capacity less than log2(2) or 1 bit. Consistent with this, the error rates become greater than zero at noise levels where our estimate of channel capacity is close to 1 bit or lower. For the unperturbed model, this occurred at noise levels of 0.5 and higher, when considering the channel capacity of the absolute response and at the noise levels of 1.0 and higher when using channel capacity of the fold change (Fig. 2a). Altogether, we predict that a population of cells can more accurately distinguish contrasting signals when relaying information via the fold change in NFκB, compared to the absolute response. This is also the case when IKKβ deactivation is disabled. Finally, this demonstrates the link between the pathway’s ability to discern different input signals is tightly related to its channel capacity.

### 3.3 Trial distributions

While the upregulation of certain surface antigens is a common feature of cancer cells, the extent of this upregulation can vary greatly across a population of cells. For example, CD19, a common marker of B cells, is present among 95% of acute lymphoblastic leukemia cells, but its abundance among myeloma cells varies between 10% and 80% depending on the patient (Nerreter et al., 2019). In order to examine how accurately CAR-4-1BB-mediated NFκB activation can relay information about encountered antigen concentration depending on the proportion of CD19-positive cells, we set out to compare the information content of the pathway when stimulated by four different antigen distributions. Each of the four distributions considered is biomodal and comprised of the same antigen-negative and antigen-positive components, but the distributions differ by the proportion of the antigen-positive component (20%, 50%, 80%, 100%) (Fig. S2).

We calculated mutual information between each antigen distribution and the NFκB activation it actuated (while all proteins of the signaling pathway were varied). Information transmitted about those four antigen distributions among a population of CAR-4-1BB cells was nearly identical for the absolute response at all noise levels (Fig. 4a). Considering the fold change in NFκB levels showed up to 1.2 bits higher mutual information between NFκB activation and the distributions with an antigen-negative component, particularly at low noise levels (Fig. 4b). In this case, information transmitted about the fully positive distribution diminished faster with increasing noise (Fig. 4b, blue), relative to distributions with an antigen-negative component. Hence, fold change in nuclear NFκB is more reflective of the encountered antigen concentrations, especially when some of the encountered targets show near-absence of antigen.

**Fig. 4:**
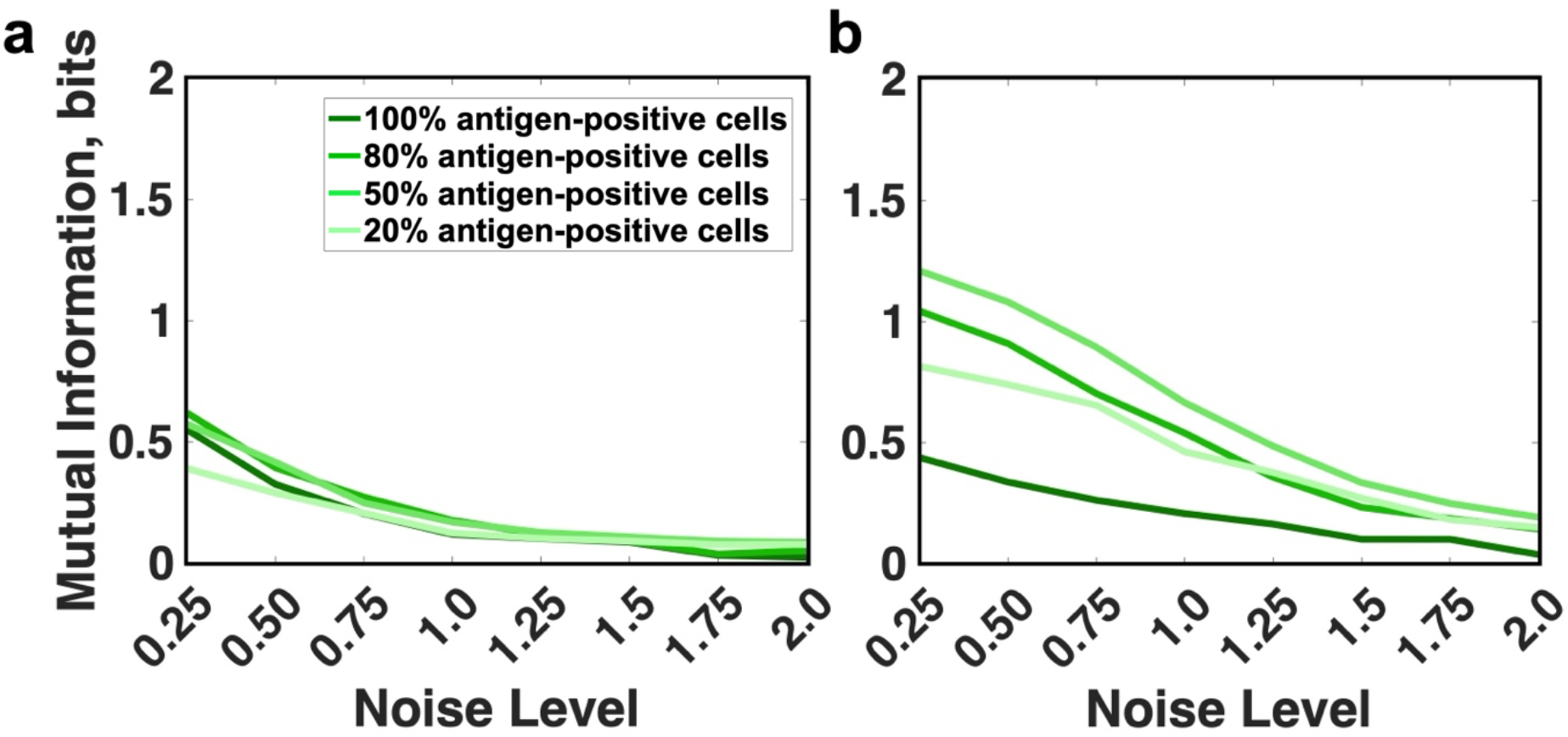
Fidelity of information transmission for different distributions of antigen concentration. (a) Mutual information between the absolute response of NFκB and the antigen; (b) Mutual information between fold increase of nuclear NFκB and the concentration of antigen (dark green, 100% antigen-positive; dark-medium, 80%; light-medium, 50%; light green, 20%).

Next, we examined same antigen distributions in the context of disabled IKKβ deactivation. Our previous work focusing on the intrinsic noise upstream of NFκB/IκBα dissociation had shown that this perturbation can increase the fidelity of upstream signaling. In line with this, mutual information between the distribution of target antigen and the distribution of absolute NFκB response it actuated increased by up to 0.5 bits when the target had an antigen-negative component (Fig. 5a compared to Fig. 4a). This improvement in information transmission was present for fully antigen-positive targets as well, albeit to a much smaller extent of up to 0.1 bits.

**Fig. 5:**
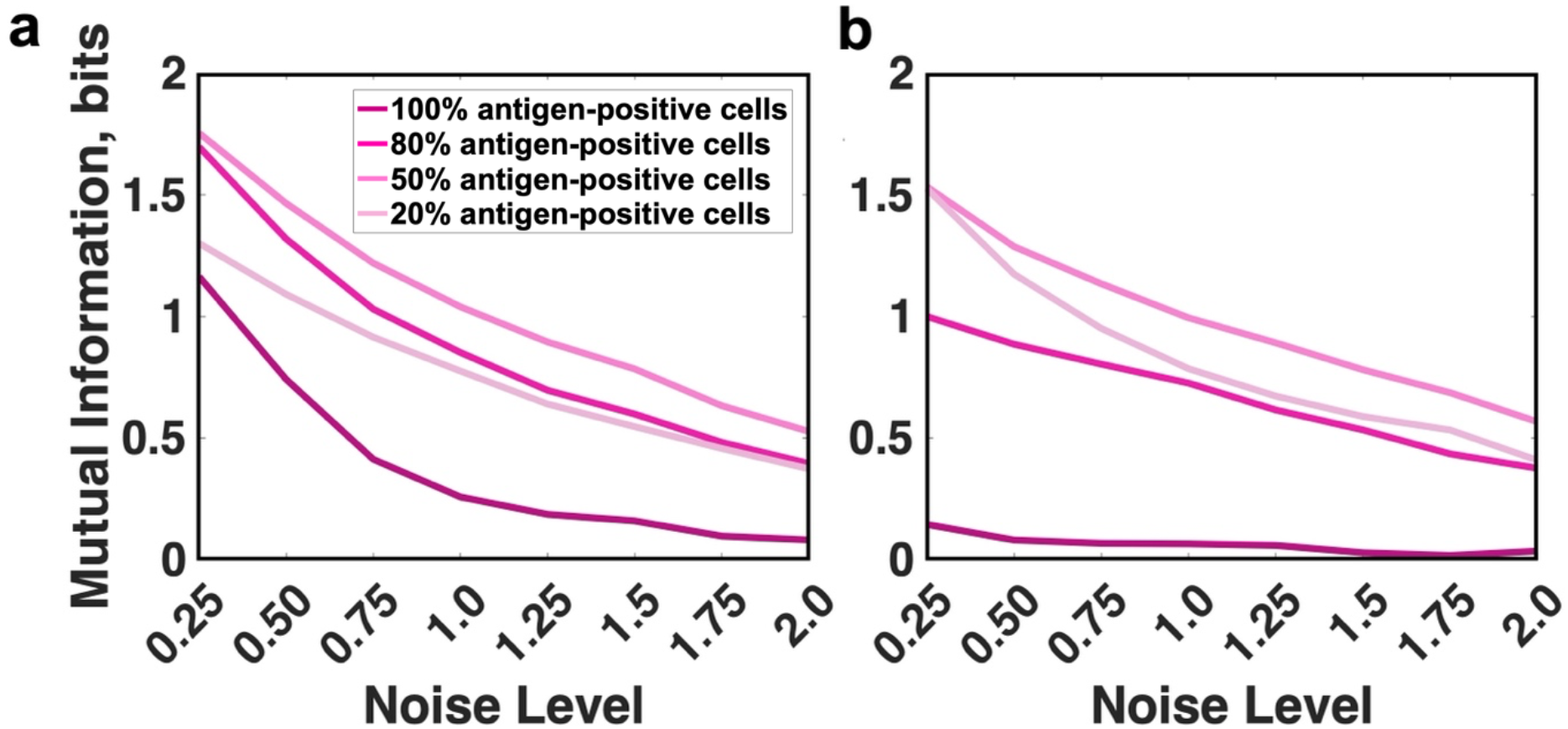
Fidelity of information transmission for different distributions of antigen concentration when deactivation of IKKβ is disabled. (a) Mutual information between the absolute response of NFκB and the antigen; (b) Mutual information between fold increase of nuclear NFκB and the concentration of antigen (dark magenta, 100% antigen-positive; dark-medium, 80%; light-medium, 50%; light magenta, 20%).

When disabling IKKβ deactivation, mutual information between antigen and NFκB fold change decreased for most cases (Fig. 5b). A notable exception was the distribution with 20% antigen positivity, which showed an increase of up 0.22 bits (Fig. 5b, light magenta). IKKβ deactivation is a mechanism to dampen the pathway response, and it is unsurprising that disabling it makes the response more accurate for an antigen distribution where the antigen-positive component has low abundance. Conversely, the fully antigen-positive distribution showed a significant deterioration of information transmission (dark magenta in Fig. 4b compared to Fig. 5b). This was in stark contrast with the case when NFκB and IκBα concentrations are fixed, where a significant increase in mutual information is observed with this perturbation. To confirm that the discrepancy is attributable to the added variability in NFκB and IκBα, we plotted the mutual information between antigen concentration and different metrics of pathway activation (Fig. S4). We observed that compared to the unperturbed system, NFκB indeed receives more information from the upstream processes. This is noted by the fact that when NFκB and IκBα were fixed, the mutual information for the perturbed system between antigen and NFκB absolute response was higher by up to 0.77 bits compared to the baseline model. However, added variability of NFκB and IκBα changed this pattern. Specifically, the mutual information between antigen and NFκB absolute response remained largely the same for the perturbed system, while the information between antigen and fold change decreased by up to 1 bit. Hence, deactivating IKKβ can have different effects on signaling accuracy depending on the prevalence of antigen in the targeted population and the response measure used to estimate information content.

In summary, we find that when all components of the signaling pathway were variable, fold change in nuclear NFκB was a more accurate measure of encountered antigen concentrations, especially if the distribution of targeted antigen contained an antigen-negative component. This finding is consistent with the greater channel capacity attributable to fold change. Perturbing the signaling pathway by disabling IKKβ deactivation resulted in an increase of mutual information between antigen concentration and the absolute response of NFκB. However, the information content of fold change mostly showed either a small decrease (for distributions with an antigen-negative component) or a drastic reduction (for distributions with only an antigen-positive component).

## 4. Discussion

In our work, we sought to develop an information-theoretic perspective of CAR-4-1BB-mediated NFκB activation based on a mathematical model of signaling. First, we estimated channel capacity of signal transmission by considering as an encoding measure either the absolute nuclear concentration of NFκB or its fold change compared to baseline levels. We found that fold change of NFκB tends to have a higher capacity for transmitting information about the encountered antigen. Furthermore, channel capacity expressed by fold change showed a weaker dependence on intrinsic noise and is close to 1 bit for the noise levels we tested. By comparing signal transmission with and without variability of IκBα and NFκB, we noted that this factor alone could diminish channel capacity by up to 1 bit, corresponding to a two-fold decrease in the number of distinct inputs the pathway accurately resolves. Next, we evaluated the NFκB pathway’s declining ability to discern contrasting signals with increasing noise levels. We found that most errors in signal transduction are attributable to a “false negative” error, whereby the pathway shows weak activation in response to a high antigen concentration. In contrast, “false positive” errors remained near zero for all noise levels tested. Similar to the case with channel capacity, cells demonstrated a better ability to discern contrasting inputs via NFκB fold change compared to the absolute response. Finally, we evaluated the ability of CAR-4-1BB-mediated NFκB signaling to accurately reflect antigen concentration for targets with different proportions of antigen-positive cells. We found that the fidelity of signal transmission for 100%, 80%, 50% and 20% antigen-positive targets is roughly identical when considering signal transmission via the absolute response of NFκB. However, when considering transmission via NFκB fold change, the fully positive distribution showed a much faster decline in transmission fidelity with increasing noise compared to distributions with antigen-negative cells.

In our previous work, we found that disabling IKKβ deactivation can greatly increase the fidelity of CAR-4-1BB-mediated NFκB activation (Tserunyan & Finley, 2022a). Here, we performed a more in-depth analysis of this perturbation. Specifically, we found that with fixed IκBα and NFκB concentrations, this manipulation could increase channel capacity by approximately 1 bit compared to the default model. This increase held in case of varying IκBα and NFκB concentrations and considering channel capacity entailed by absolute response. However, the increase in capacity entailed by fold change was modest when varying IκBα and NFκB concentrations. Additionally, we found that the perturbation improved the pathway’s ability to discern contrasting signals with a 20-25% lower error rate. Finally, we observed that disabling IKKβ deactivation could improve the fidelity of signal transmission for input distributions with an antigen-negative component if we consider encoding via absolute response. Interestingly, in the case of signal encoding via fold change, the perturbation drastically reduced signaling fidelity for the fully antigen-positive distribution.

Previous studies of a signaling pathways have demonstrated that channel capacities of most pathways are within 0.5-1.75 bits (Topolewski & Komorowski, 2021). An experimental estimate of the channel capacity of NFκB signaling in TNF-stimulated fibroblasts suggested the value of 0.92 bits, implying that the pathway can resolve only 2^0.92^ ≈ 2 input concentrations (Tudelska et al., 2017). However, pathways can increase their information content by encoding stimulus strength not in the absolute concentration of the secondary messenger, but in the ensemble of multiple cross-wired effectors or in the fold change of the messenger’s concentration (Suderman et al., 2017; Topolewski & Komorowski, 2021). For example, some experimental findings suggest that NFκB fold change is more informative than its absolute abundance (R. E. C. Lee et al., 2014). This is consistent with our finding that under most simulation scenarios, fold change shows a higher channel capacity. Prior modeling studies have found that a relatively small change in channel capacity from 0.85 bits to 0.6 bits can increase the number of incorrectly responding cells by twofold (Tabbaa & Jayaprakash, 2014). Thus, enhancing the fidelity of signaling pathways that actuate the therapeutic function of CAR T cells could result in a better alignment between the response of the CAR cell and the target it encounters. Given our finding that the same perturbation to the signaling pathway affects information transmission differently depending on the presence of antigen-negative targets, a successful engineering approach would consider features of likely targets for the more precise design of CAR T cells.

Along with the novel findings in our work, we acknowledge certain areas that could be improved upon in the future. One such area is the correspondence between our model and the experimental system. Due to the scarcity of available measurements, we had to make order-of-magnitude estimates for some parameters by knowing the timescale at which the underlying process proceeds. Additionally, other parameter values were based on studies of NFκB activation in fibroblasts via the TNF receptor (TNFR). The basis for applying these data on CAR-4-1BB-mediated NFκB activation is that 4-1BB and TNFR belong to the same superfamily of receptors and trigger cell signaling via a similar mechanism of recruiting TRAF proteins (Zapata et al., 2018). It is possible that more numerically accurate findings may be produced if our model is calibrated against measurements performed in CAR-4-1BB cells. Another limitation that we recognize is the assumption that all proteins in a heterogeneous population are distributed according to a lognormal distribution with the same scale parameter. Lognormality of protein concentrations has been widely recognized for steadily growing cells, while experimental measurements have found that the ratio of standard deviation to the mean is constant (Furusawa et al., 2005). By our calculations, this ratio implies an identical scale parameter for protein distributions, roughly equal to 0.5. Notably, at a noise level of 0.5, our simulations predict a channel capacity of 0.85 bits for the absolute response of NFκB, while the experimentally measured value is 0.92 bits. Nevertheless, both lognormality and identical scale parameters for protein distribution depend on the steady growth of cells. In cases when this assumption does not hold, different proteins could have different variability levels potentially causing unexpected behaviors of the system. This is underscored by the fact that disabling IKKβ deactivation had diametrically opposing consequences on the accuracy of signal transmission depending on the variability of IκBα and NFκB. We can update the model as more information becomes available regarding the distributions of the concentration of signaling species.

In conclusion, we aimed to use mathematical modeling and information theory to grasp CAR-4-1BB-mediated NFκB activation. We first estimated the channel capacity of this transduction pathway under different conditions. Then, we examined the pathway’s ability to distinguish low and high signals in the presence of intrinsic noise. Finally, we evaluated how accurately NFκB activation reflects encountered antigen concentration in a tumor population. Our findings suggest that fold change in the nuclear concentration of NFκB has a higher channel capacity than the absolute response. We also found that due to intrinsic noise, the response of the pathway tends to underestimate the concentration of target antigen. Finally, we discovered that disabling IKKβ deactivation could improve signaling fidelity when encountering targets that have antigen-negative cells. Thus, our analysis offers new insights into biological signaling and can inform cell engineering practices.

## Data availability

All datasets and scripts for this research article are available via the following link: https://github.com/FinleyLabUSC/InfoTheory-NFkB.

## Author Contributions

SDF conceptualized and directed the project. VT performed relevant simulations and developed figures. VT wrote the draft manuscript. SDF and VT jointly reviewed and edited the manuscript. SDF provided funding for the project. All authors have read and approved the final manuscript.

## Acknowledgements

The authors would like to acknowledge the constructive critiques and suggestions from the members of the Computational Systems Biology group at the University of Southern California.

