## Supplementary Figures for "Information-theoretic analysis of a model of CAR-4-1BB-mediated NFκB activation"

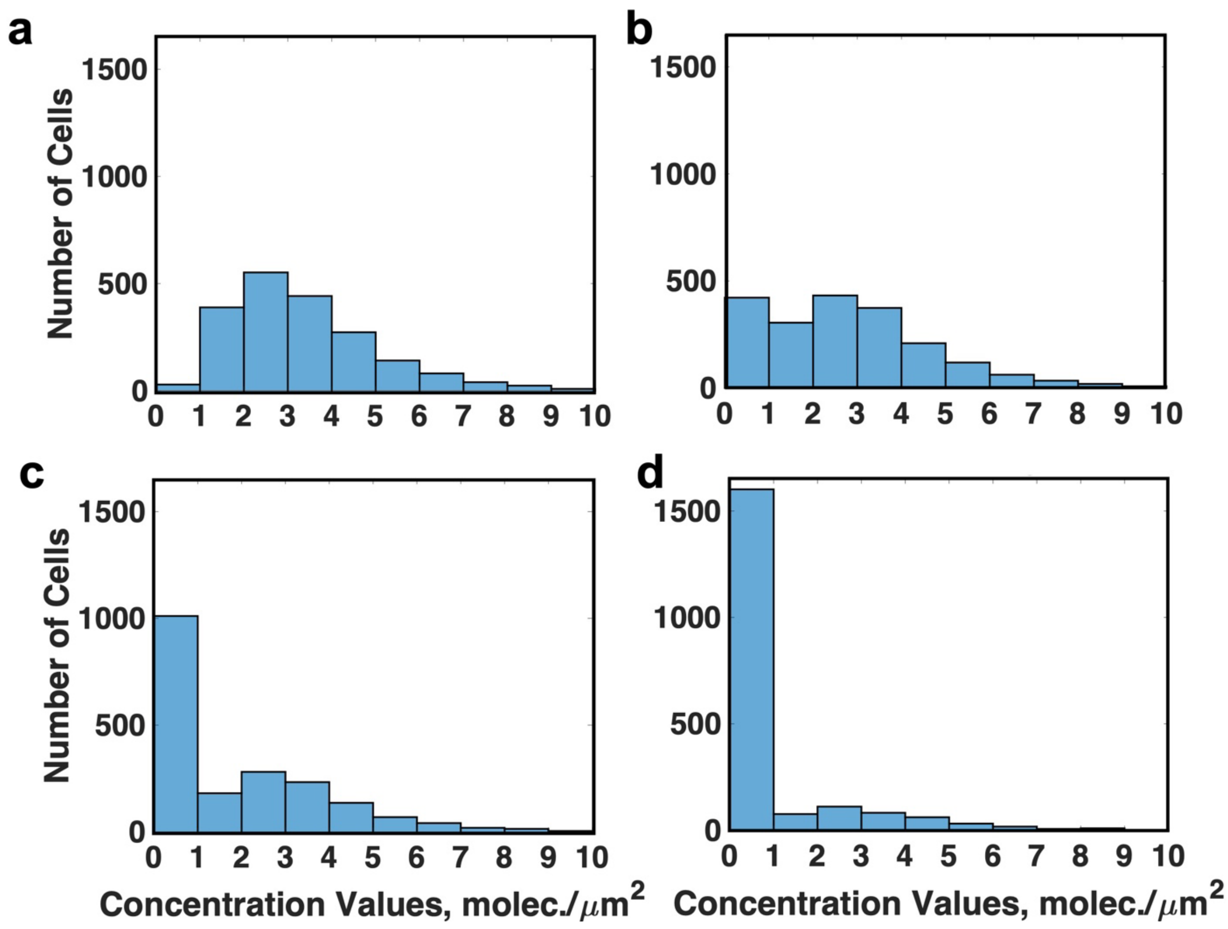


**Fig. S1**: Different distributions of antigen concentration for probing the fidelity of CAR-4-1BB-mediated NFκB activation, with identical positive and negative components in varying proportions. (a) 100% cells antigen-positive; (b) 80%; (c) 50%; (d) 20%.

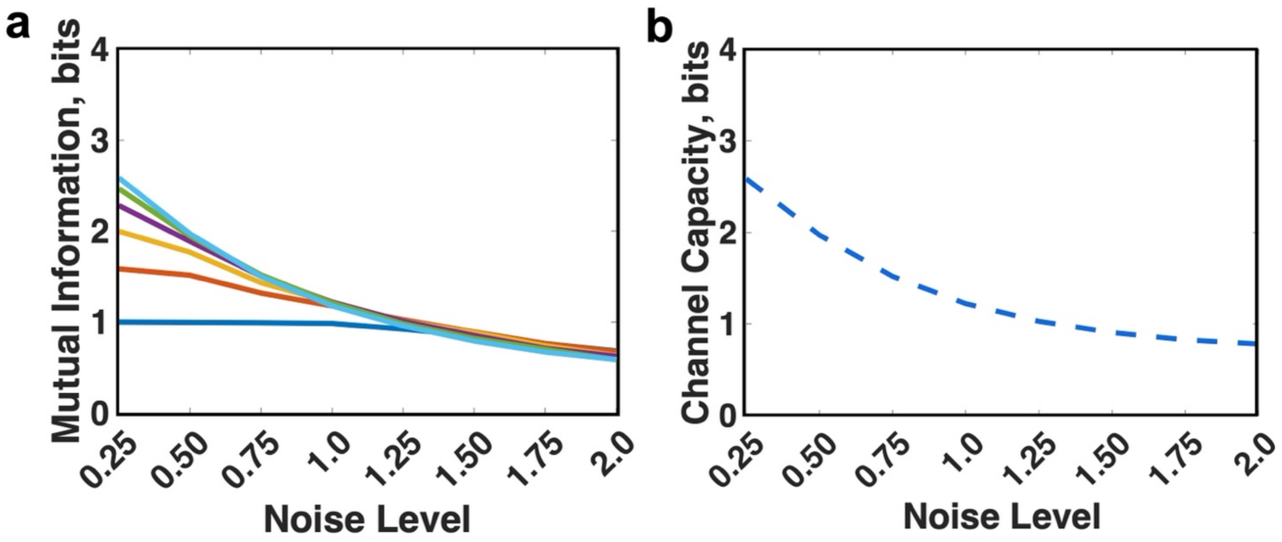


**Fig. S2**: Example of computing channel capacity. (a) Mutual information between different candidates for a capacity-maximizing antigen distribution and the pathway response (blue, two antigen concentrations; orange, three; yellow, four; purple, five; green, six; cyan, seven). (b) Channel capacity estimated as the maximum mutual information achievable at each noise level, based on A (blue curve identical to that in **Fig. 2a**).

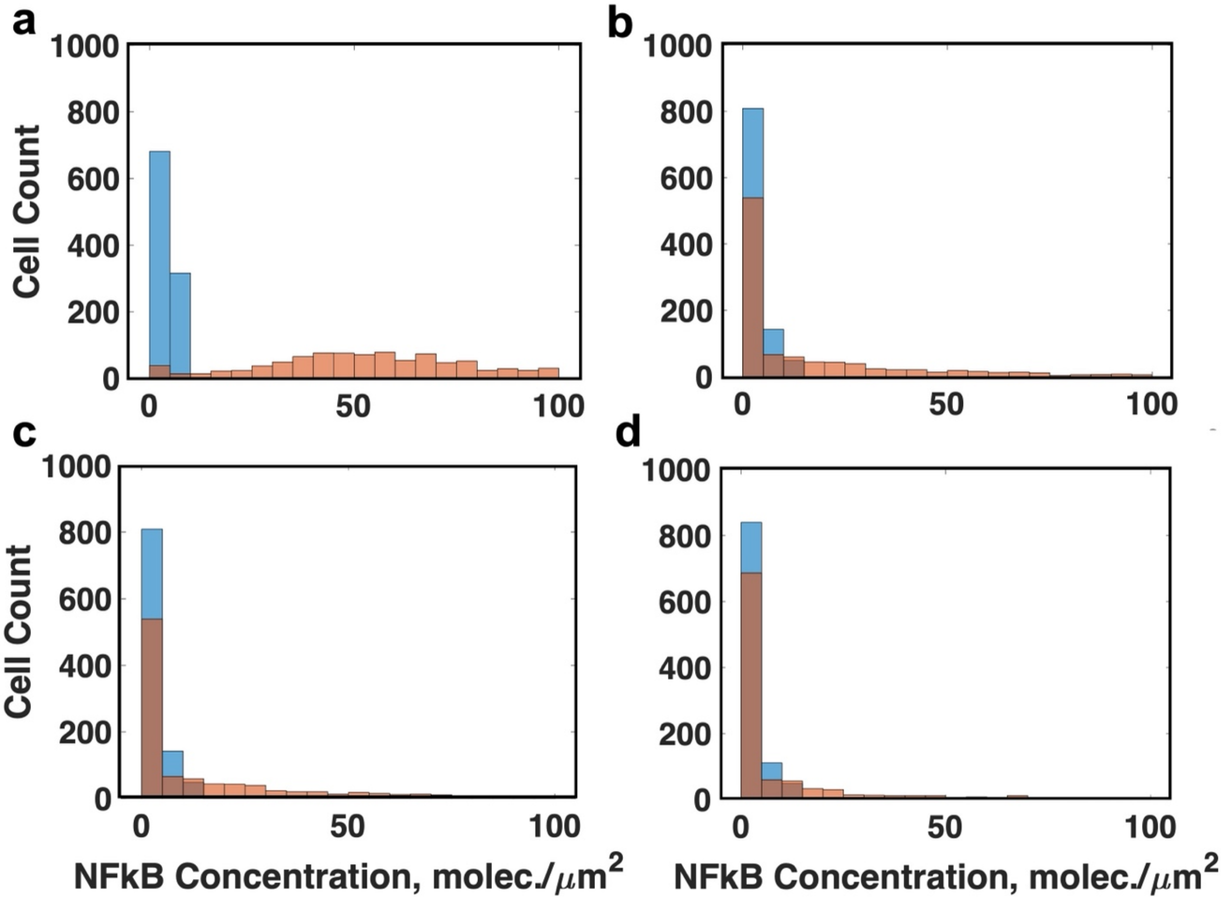


**Fig. S3**: Deteriorating ability to discern contrasting signals with increasing intrinsic noise. Noise level set at (a) 0.5, (b) 1.0, (c) 1.5, (d) 2.0. Blue shows the unperturbed pathway stimulated with antigen concentration of 0.7 molecules/ $\mu m^2$, orange shows results from stimulation with antigen concentration of 70 molecules/$\mu m^2$. Note that with increasing noise levels, the response distribution of "high" stimulation becomes closer to that of the "low" stimulation. For this reason, most of the response values of "high" stimulation become more characteristic of "low"-stimulated cells.

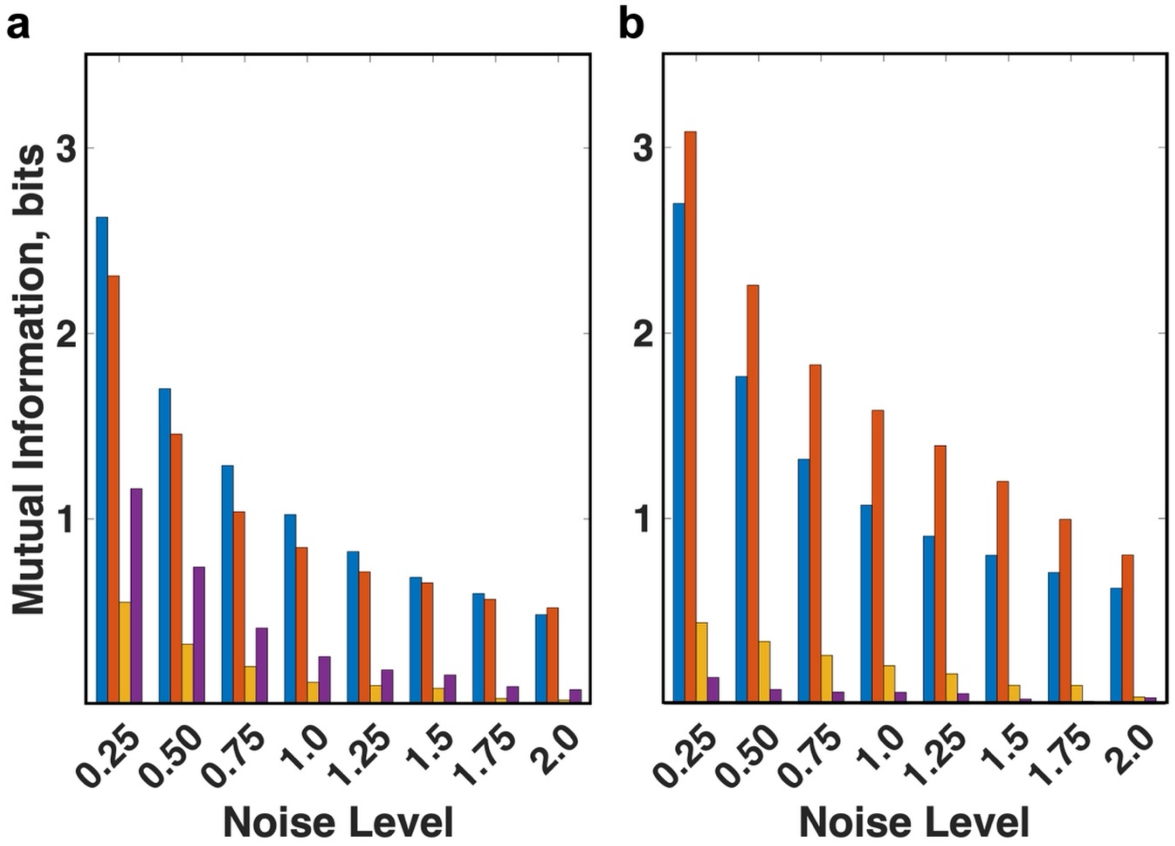


**Fig. S4**: Mutual information between 100% antigen-positive distribution and various metrics of pathway activation for (a) the unperturbed system and (b) with disabled deactivation of IKKβ. Mutual information between antigen concentration and enzymatically active IKKβ in blue, absolute response of NFκB in orange (if IκBα and NFκB are fixed), absolute response of NFκB in yellow (if IκBα and NFκB are variable), fold change in nuclear NFκB in purple (if IκBα and NFκB are variable).
